# Bioshake: a Haskell EDSL for bioinformatics pipelines

**DOI:** 10.1101/529479

**Authors:** Justin Bedő

## Abstract

**Background:** Typical bioinformatics analysis comprise long running computational pipelines. An important part of producing reproducible research is the management and execution of these computational pipelines to allow robust execution and to minimise errors. Bioshake is an embedded domain specific language embedded in Haskell for specifying and executing computational pipelines in bioinformatics that significantly reduces the possibility of errors occurring.

**Results:** Unlike other pipeline frameworks, Bioshake raises many properties to the type level to allow the correctness of a pipeline to be statically checked during compilation, catching errors before any lengthy execution process. Bioshake builds on the Shake build tool to provide robust dependency tracking, parallel execution, reporting, and resumption capabilities. Finally, Bioshake abstracts execution so that jobs can either be executed directly or submitted to a cluster.

**Conclusions:** Bioshake is available at http://github.com/papenfusslab/bioshake.

## 1 Background

Bioinformatics pipelines are typically composed of numerous programs and stages coupled together loosely using intermediate files. These pipelines tend to be quite complex and require much computational time, hence a good pipeline must be able to manage intermediate files, guarantee rentrability – the ability to re-enter a partially run pipeline and continue from the latest point – and also provide methods to easily describe pipelines.

We present *bioshake*: a Haskell Embedded Domain Specific Language (EDSL) for bioinformatics pipelines. The use of a language with strong types gives our framework several advantages over existing frameworks [Leipzig, 2016, Goodstadt, 2010, Amstutz et al., 2016, wdl, 2012, Vivian et al., 2017]:

1. The type system is strongly leveraged to prevent errors in the pipeline construction during compilation. Errors such as mismatching file types, combining samples mapped against different references, or failing to sort a Sequence Alignment Map (SAM) file before a stage that requires sorting all result in a compile error rather than a runtime error. This catches errors significantly earlier, reducing debugging time. As bioinformatics pipelines tend to have long runtimes, this is especially advantageous. To the best of our knowledge, this is the first bioinformatics pipeline framework to use strong typing and type inference to prevent specification errors during compile time.
2. Naming of outputs at various stages of a pipeline are abstracted by bioshake. Output at a stage can be explicitly named if they are desired outputs. Thus, the burden of constructing names for temporary files is alleviated. This is similar in spirit to Sadedin et al. [2012] who also allow abstraction away from explicit filenames.
3. Bioshake builds on top of *Shake*, an industrial strength build tool also implemented as an EDSL in Haskell. Bioshake thus inherits the reporting features, robust dependency tracking, and resumption capabilities offered by the underlying Shake architecture.
4. Unlike underlying shake that expects dependencies to be specified (i.e., in a DAG the arrows point from the target back towards the source(s)), bioshake allows forward specification of pipelines (i.e., the arrows point forward). As bioinformatics pipelines tend to be quite long and mostly linear, this eases the cognitive burden during pipeline design and also improves readability.
5. Non-linear pipelines are constructed using typical Haskell constructs such as maps and folds. Combinators are available for the most common grouping of outputs together for a subsequent stage. However, as the main data type is recursively defined, outputs of a stage can always be referenced by subsequent stages without explicit non-linear constructs (i.e., the alignments used for variant calling are available for a subsequent variant annotation stage without explicitly introducing non-linearity).

Bioshake in essence is an EDSL for specifying pipelines that compiles down to an execution engine (shake). In this respect, it is similar to other specification languages such as Common Workflow Language (CWL) [Amstutz et al., 2016] and Workflow Description Language (WDL) [wdl, 2012], but executes on top of shake. Table 1 provides a high level feature overview of Bioshake when compared to several other pipeline specification language, pipeline EDSLs, and execution engines. We will further elaborate on the unique features of Bioshake:

**Table 1:** High level feature comparison of Bioshake with other execution engines (Toil, Cromwell), specification languages (WDL, CWL), and EDSLs (Ruffus). Dashes indicate that feature is not applicable.

|  | Ruffus | Toil | Cromwell | WDL | CWL | Bioshake |
| --- | --- | --- | --- | --- | --- | --- |
| Embedded DSL | ✓ | — | — |  |  | ✓ |
| Python | ✓ | ✓ | — |  |  |  |
| Strong static typing |  |  | — |  |  | ✓ |
| Type inferencing |  |  | — |  |  | ✓ |
| Extrinsic specification |  |  | — |  |  | ✓ |
| Intrinsic specification | ✓ | ✓ | — | ✓ | ✓ | ✓ |
| Functional language |  |  | — |  |  | ✓ |
| Container integration |  | ✓ | ✓ | — | — |  |
| Cloud computing integration |  | ✓ | ✓ | — | — |  |
| Cluster integration (Torque) | — | ✓ | ✓ | — | — | ✓ |
| Cluster integration (Slurm) | — | ✓ | ✓ | — | — |  |
| Cluster integration (SGE) | — | ✓ | ✓ | — | — |  |
| Cluster integration (LSF) | — | ✓ |  | — | — |  |
| Cluster integration (DRMAA) | ✓ |  |  | — | — |  |
| Direct execution | ✓ | ✓ | ✓ | — | — | ✓ |

### Strong type-checking

The use of a language with strong types gives our framework several advantages over existing frameworks [Leipzig, 2016, Goodstadt, 2010, Sadedin et al., 2012, Amstutz et al., 2016, wdl, 2012, Vivian et al., 2017]. Our framework leverages Haskell’s strong type-checker to prevent many errors that can arise in the specification of a pipeline. As an example, file formats are statically checked by the type system to prevent specification of pipelines with incompatible intermediate file formats. Furthermore, tags are implemented through Haskell type-classes to allow metadata tagging, allowing various properties of files – such as whether a bed file is sorted – to be statically checked. Thus, a miss-specified pipeline will simply fail to compile, catching these bugs well before the lengthy execution. This feature is not present in other bioinformatics pipeline frameworks such as those reviewed by Leipzig [2016].

### Intrinsic and extrinsic building

Our framework builds upon the Shake EDSL [Mitchell, 2012], which is a make-like build tool. Similarly to make, dependencies in shake are specified in an *extrinsic* manner (called internal/external by Leipzig [2016]), that is a build rule will define its input dependencies based on the output file path. Our EDSL compiles down to shake rules, but allows the specification of pipelines in an *intrinsic* fashion, whereby the processing chain is explicitly stated and hence no filename based dependency graph needs to be specified. However, as bioshake compiles to shake, both extrinsic and intrinsic rules can be mixed, allowing a choice to be make to maximise pipeline specification clarity. For example, small “side” processing like generation of indices can be specified extrinsically, removing the need for an explicit index step in the pipeline specification.

Furthermore, the use of explicit sequencing for defining pipelines allows abstraction away from the filename level: intermediate files can be automatically named and managed by bioshake, removing the burden of naming the intermediate files, with only desired outputs requiring explicit naming.

**Example 1** The following is an example of a pipeline expressed in the bioshake EDSL:

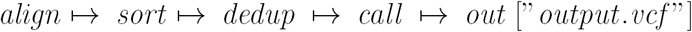

From this example it is clear what the *stages* are, and the names of the files flowing between stages is implicit and managed by Bioshake. The exception is the explicitly named output, which is the output of the whole pipeline. Note that non-linearity is handled by constructors that accept the extra inputs, but pipelines can always recurse backwards along ↦ to retrieve prior build products (e.g., to fetch Binary Alignment Map (BAM) files used to generate a set of variant calls), reducing the need for non-linearity.

### Extends a robust build system

Finally, the Bioshake EDSL compiles to Shake [Mitchell, 2012], an industrial strength build tool also implemented as an EDSL in Haskell. Bioshake thus inherits the reporting features, robust dependency tracking, and resumption capabilities offered by the underlying Shake framework. Though Bioshake is not the first EDSL for bioinformatics pipelines [Goodstadt, 2010, Leipzig, 2016], to the best of our knowledge it is the first EDSL in Haskell and the first to use a deep type embedding to prevent invalid pipeline specifications.

## 2 Implementation

### 2.1 Core data types

Bioshake is build using a tagless-final style [Carette et al., 2009] around the following datatype:

~~~
**data** *a* ↦ *b*
  **where**
    (↦) ∷ *a* → *b* → *a* → *b*
**infixl** 1 ↦
~~~

This datatype represents the conjunction of two stages *a* and *b*. As we are compiling to shake rules, the *Buildable* class represents a way to build thing of type *a* by producing shake actions:

~~~
**class** *Buildable a*
  **where**
   *build* ∷ *a* → *Action* ()
~~~

Finally, as we are ultimately building files on disk, we use a typeclass to represent types that can be mapped to filenames:

~~~
**class** *Pathable a*
  **where**
   *paths* ∷ *a* → [*FilePath*]
~~~

### 2.2 Defining stages

A stage – for example *align*ing and *sort*ing – is a type in this representation. Such a type is an instance of *Pathable* as outputs from the stage are files, and also *Buildable* as the stage is associated with some shake actions required to build the outputs. We give a simple example of declaring a stage that sorts bam files.

**Example 2** Consider the stage of sorting a bed file using samtools. We first define a datatype to represent the sorting stage and to carry all configuration options needed to perform the sort:

~~~
*data Sort = Sort*
~~~

This datatype must be an instance of *Pathable* to define the filenames output from the stage. Naming can take place according to several schemes, but here we will opt to use hashes to name output files. This ensure the filename is unique and relatively short.

~~~
**instance** *Pathable a* ⟹ *Pathable (a* ↦ *Sort)*
  **where**
  *paths* (a ↦ _) = **let**
                    *inputs = paths a*
                  **in**
                  [*hash inputs* + + ”:*sort:bed* ”]
~~~

In the above, *hash* ∷ *Binary a ⇒ a* → *String* is a cryptographic hash function such as sha1 with base32 encoding. Many choices are appropriate here.

Finally, we describe how to sort files by making *Sort* an instance of *Buildable*:

~~~
**instance** (*Pathable a, IsBam a*) ⇒*Buildable* (*a* ↦*Sort*)
  **where**
    *build p*@(*a* ↦ _) = **let**
                         [*input*] = *paths a*
                         [*out*] = *paths p*
                       **in**
                       *cmd* ”*samtools sort* ” [*input*] [” − *o*”,* out*]
~~~

Note here that *IsBam* is a precondition for the instance: the sort stage is only applicable to BAM files. Likewise, the output of the sort is also a BAM file, so we declare that too:

~~~
**instance** *IsBam* (*a* ↦ *Sort*)
~~~

The tag *IsBam* itself can be declared as the empty typeclass *class IsBam a*. See section 2.4 for a discussion of tags and their utility.

### 2.3 Compiling to shake rules

The pipelines as specified by the core data types are compiled to shake rules, with shake executing the build process. The distinction between *Buildable* and *Compilable* types are that the former generate shake *Action*s and the latter shake *Rules*. The *Compiler* therefore extends the *Rules* monad, augmenting it with some additional state:

~~~
**type** *Compiler* = *StateT* (*S.Set* [*FilePath*]) *Rules*
~~~

The state here captures rules we have already compiled. As the same stages may be applied in several concurrent pipelines (i.e., the same preprocessing may be applied but different subsequent processing defined) the set of rules already compiled must be maintained. When compiling a rule, the state is checked to ensure the rule is new, and skipped otherwise. The rule compiler evaluates the state transformer, initialising the state to the empty set:

~~~
*compileRules* ∷ *Compiler* ()→ *Rules* ()
*compileRules p* = *evalStateT p mempty*
~~~

A compilable typeclass abstracts over types that can be compiled:

~~~
**class** *Compilable a*
**where**
*compile* ∷ *a → Compiler* ()
~~~

*a* ↦ *b* is *Compilable* if the input and output paths are defined, the subsequent stage *a* is *Compilable*, and *a* ↦ *b* is *Buildable*. Compilation in this case defines a rule to build the output paths with established dependencies on the input paths using the *build* function. These rules are only compiled if they do not already exist:

~~~
*instance* (*Pathable a, Pathable* (*a ↦ b*),* Compilable a, Buildable* (*a ↦ b*)) *⇒ Compilable* (*a ↦ b*)
 **where**
  *compile pipe*(*a* ↦*b*) = *do
    let outs* = *paths pipe
    set* ←* get
    when* (*outs* ‘*S.notMember*‘ *set*) $ *do
       lift* $ *outs* &%*>* _ *do
         need* (*paths a*)
         *build pipe
     put* (*outs* ‘*S.insert*‘ *set*)
  *compile a*
~~~

### 2.4 Tags

Bioshake uses tags to ensure type errors will be raised if stages are incompatible. We have already seen in example 2 the use of *IsBam* to ensure the input file format of *Sort* is compatible. By convention, Bioshake uses the file extension prefixed by *Is* as tags for filetype, e.g.,: *IsBam, IsSam, IsVCF*.

Other types of metadata are used such as if a file is sorted (*Sorted*) or if duplicate reads have been removed (*DeDuped*) or marked (*DupsMarked*). These tags allow input requirements of sorting or deduplication to be captured when defining stages. Properties, where appropriate, can also automatically propagate down the pipeline; for example, once a file is *DeDuped* all subsequent outputs carry the *DeDuped* tag:

~~~
 **instance** *Deduped a ⇒ Deduped* (*a ↦ b*)
~~~

Finally, the tags discussed so far have been empty type classes, however tags can easily carry more information. For example, bioshake uses a *Referenced* tag to represent the association of a reference genome. This tag is defined as

~~~
 **class** *Referenced*
  **where**
   *getRef* ∷ *FilePath*
 **instance** *Referenced a* ⇒ *Referenced* (*a* ↦ *b*)
~~~

This tag allows stages to extract the path to the reference genome and automatically propagates down the pipeline allowing identification of the reference at any stage.

### 2.5 EDAM ontology

EDAM [Ison et al., 2013] is an ontology containing terms and concepts that are prevalent in the field of bioinformatics. As it is a formal ontology, the terms are organised into a hierarchical tree structure, with each term containing reference to parent terms. EDAM can be used with the flat tagging structure introduced in the previous section through the use of template Haskell to establish the tree.

Bioshake provides the EDAM ontology in the EDAM module. This module provides EDAM terms identified by their short name, along with some template Haskell for associating EDAM terms to types. For example, the *FASTQ-illumina* term (http://edamontology.org/format_1931) is represented by the tag *FastqIllumina* and a type can be tagged using the *is* template Haskell function, for example:

~~~
 **import** *Bioshake.EDAM*
 **data** *MyType* = *MyType*
 $(*is ″MyType ″FastqIllumina*)
~~~

Output of stages (e.g., types of *a*↦ *MyType*) can equally be tagged using the *allP* template Haskell function:

~~~
 $(*isP ″MyType ″FastqIllumina*)
~~~

These template Haskell functions declare the given type to be instances of all parents of the EDAM term, allowing tag matching at any level in the hierarchy. These EDAM types can be used similarly to tags as described in section 2.4.

### 2.6 Abstracting the execution platform

In example 2, the shake function *cmd* is directly used to execute samtools and perform the build, however it is useful to abstract away from *cmd* directly to allow the command to be executed instead on (say) a cluster, cloud service, or remote machine. Bioshake achieves this flexibility by using free monad transformers to provide a function *run* – the equivalent of *cmd* – but where the actual execution may take place via submitting a script to a cluster queue, for example.

To this end, the datatype for stages in bioshake are augmented by a free parameter to carry implementation specific default configuration – e.g., cluster job submission resources. In the running example of sorting a bed file, the augmented datatype is **data** *Sort c* = *Sort c*.

### 2.7 Reducing boilerplate

Much of the code necessary for defining a new stage can be automatically written using template Haskell. This allows very succinct definitions of stages increasing clarity of code and reducing boilerplate. Bioshake has template Haskell functions for generating instances of *Pathable* and *Buildable*, and for managing the tags.

**Example 3** Example 2 can be simplified by using template Haskell considerably. First we have the augmented type definitions:

~~~
 **data** *Sort c* = *Sort c*
~~~

The instances for *Pathable* and the various tags can be generated with the template Haskell splice

~~~
 $(*makeTypes ″Sort* [*″IsBam, ″Sorted*] [])
~~~

This splice generates a *Pathable* instance using the hashed path names, and also declares the output to be instances of *IsBam* and *Sorted*. The first tag in the list of output tags determines the file extension. The second empty list allows the definition of *transient* tags; that is the tags that if present on the input paths will hold for the output files after the stage. Finally, given a generic definition of the build

~~~
 *buildSort t* _ (*paths* → [*input*]) [*out*] =
    *run* ”*samtools sort* ” [*input*] [”-@”,* show t*] [”-o”,* out*]
~~~

the *Buildable* instances can be generated with the splice

~~~
 $(*makeThreaded ″Sort* [*″IsBam*] *fbuildSortBam*)
~~~

This splice takes the type, a list of required tags for the input, and the build function. Here, the build function is passed the number of threads to use, the *Sort* object, the input object and a list of output paths.

## 3 Results and Discussion

We have presented a framework for describing and executing bioinformatics pipelines. The framework is an EDSL in Haskell and built on shake. This allows us to leverage the robustness of shake, and also the power of Haskell’s type system to prevent many types of errors in pipeline construction. This is of great benefit for bioinformatics pipelines, as they tend to be long running and thus catching errors during compile reduces the debugging time significantly.

Though this library is built around Shake as the execution engine, the core value lies in the unique abstraction and use of types to capture metadata. It is feasible to compile a specification to a different backend instead of Shake, such as Toil [Vivian et al., 2017] or Cromwell [cro, 2015] via CWL [Amstutz et al., 2016] or WDL [wdl, 2012]. This would allow leveraging of the cloud and containerisation facilities of Toil and Cromwell. The abstraction used may also be useful in other domains where long data-transformation stages are applied, such as data mining on large datasets.

Though many errors are currently caught by the type system, there are still classes of errors that are not. Notably, the Pathable class instance maps stages to lists of files with unknown length. Thus, the number of files expected to be exchanged between two stages may differ, causing a runtime error. This could in principle be caught by using lists of typed length, however this would increase the complexity for users. Bioshake attempts to strike a balance between usability and type safe guarantees.

## 4 Conclusions

We have presented a unique EDSL in Haskell for specifying bioinformatics pipelines. The Haskell type checker is used extensively to prevent specification errors, allowing many errors to be caught during compilation rather than runtime. To our knowledge, this is the first bioinformatics pipeline framework in Haskell, as well as the first formalisation of bioinformatics pipelines and their attributes in a type system from the Hindley–Milner family.

## 5 Availability and Requirements

**Project name:** bioshake

**Project home page:** http://github.com/papenfusslab/bioshake

**Operating system(s)**: Windows, Linux, MacOS X

**Programming language:** Haskell

**License:** ISC

## Abbreviations

BAM: Binary Alignment Map
CWL: Common Workflow Language
DRMAA: Distributed Resource Management Application API [Troger et al., 2007]
EDSL: Embedded Domain Specific Language
SAM: Sequence Alignment Map

## Declarations

### Ethics approval and consent to participate

not applicable.

### Consent for publication

not applicable.

### Availability of data and material

bioshake is available at http://github.com/papenfusslab/bioshake.

### Competing interests

the authors declare that they have no competing interests.

### Funding

JB is supported by the Stafford Fox Centenary Fellowship in Rare Cancer.

### Authors’ contributions

JB designed the framework, implement the design, and wrote the manuscript.

## Acknowledgements

I thank Tony Papenfuss for supporting this work and helpful discussions. I also thank Leon di Stefano and Jan Schröder for helpful discussions.

